# Human disease-specific cell signatures in non-lesional tissue in Multiple Sclerosis detected by single-cell and spatial transcriptomics

**DOI:** 10.1101/2023.12.20.572491

**Authors:** Matti Lam, Dylan Lee, Ivy Kosater, Anthony Khairallah, Mariko Taga, Ya Zhang, Masashi Fujita, Sukriti Nag, David A. Bennett, Philip De Jager, Vilas Menon

## Abstract

Recent investigations of cell type changes in Multiple Sclerosis (MS) using single-cell profiling methods have focused on active lesional and peri-lesional brain tissue, and have implicated a number of peripheral and central nervous system cell types. However, an important question is the extent to which so-called “normal-appearing” non-lesional tissue in individuals with MS accumulate changes over the lifespan. Here, we compared post-mortem non-lesional brain tissue from donors with a pathological or clinical diagnosis of MS from the Religious Orders Study or Rush Memory and Aging Project (ROSMAP) cohorts to age– and sex-matched brains from persons without MS (controls). We profiled three brain regions using single-nucleus RNA-seq: dorsolateral prefrontal cortex (DLPFC), normal appearing white matter (NAWM) and the pulvinar in thalamus (PULV), from 15 control individuals, 8 individuals with MS, and 5 individuals with other detrimental pathologies accompanied by demyelination, resulting in a total of 78 samples. We identified region– and cell type-specific differences in non-lesional samples from individuals diagnosed with MS and/or exhibiting secondary demyelination with other neurological conditions, as compared to control donors. These differences included lower proportions of oligodendrocytes with expression of myelination related genes MOBP, MBP, PLP1, as well as higher proportions of CRYAB+ oligodendrocytes in all three brain regions. Among microglial signatures, we identified subgroups that were higher in both demyelination (TMEM163+/ERC2+), as well as those that were specifically higher in MS donors (HIF1A+/SPP1+) and specifically in donors with secondary demyelination (SOCS6+/MYO1E+), in both white and grey matter. To validate our findings, we generated Visium spatial transcriptomics data on matched tissue from 13 donors, and recapitulated our observations of gene expression differences in oligodendrocytes and microglia. Finally, we show that some of the differences observed between control and MS donors in NAWM mirror those previously reported between control WM and active lesions in MS donors. Overall, our investigation sheds additional light on cell type– and disease-specific differences present even in non-lesional white and grey matter tissue, highlighting widespread cellular signatures that may be associated with downstream pathological changes.

## Introduction

Multiple sclerosis (MS) is a neurological disorder characterized by the presence of perivascular brain lesions associated with infiltration of peripheral immune cells and subsequent demyelination, neuronal injury, and accelerated brain atrophy. Unlike other neurodegenerative diseases, this brain atrophy is not regional; it is global and involves both the cortex and deep gray matter such as the thalamus^1^ The disease itself can manifest in heterogeneous symptomologies, which include relapsing-remitting (RRMS), primary progressive, and secondary progressive forms^2–4^. However, a common theme across these forms is damage to oligodendrocytes, demyelination, and ultimately regional atrophy. Recent molecular studies have focused on characterizing the cellular and spatial composition of lesion centers and peri-lesional areas^5–7^, identifying a host of resident glial and infiltrating immune cell types with altered gene expression signatures. However, an open question is how tissue without active lesions is affected long-term in individuals with MS. In particular, grey matter changes have been observed in brain regions that typically do not exhibit large numbers of acute lesions^1^, including cortical gray matter and the thalamus. Here, we sought to identify cellular and molecular changes in these tissues as well as in non-lesional white matter, with the goal of identifying potentially targetable cell types associated with systemic effects of acute MS lesions, including changes in the thalamus and cortical grey matter.

## Results

### Identifying and localizing broad cell classes and subtypes

We selected 28 donors from the ROSMAP^8^ cohort (Fig 1A); they included the following 3 groups: (i) individuals with either a clinical or pathologic diagnosis of MS (MS Group, n=8), (ii) individuals with demyelination observed post-mortem, but no MS diagnosis or lesions (Secondary demyelination, n=4 with a pathologic diagnosis of Alzheimer’s Disease (AD), n=1 with stroke), and (iii) reference participants with with no ante-mortem cognitive or motor symptoms, and no evidence of demyelination post-mortem (Control, n=15). Importantly, individuals in the Control group were selected to have the same broad distribution of sex and Alzheimer’s Disease pathology (neurofibrillary tangles and amyloid plaques) as those in the two demyelination groups (Fig 1B). Crucially, our focus is not to replicate previous studies that have looked at cell type signatures in acute lesions, but to examine systemic changes present in non-lesional brain tissue. To this end, we profiled three brain regions from each individual, the dorsolateral prefrontal cortex (DLPFC), the normal appearing white matter from anterior watershed (NAWM) and the pulvinar in the thalamus (Pulvinar) using single nucleus RNA sequencing (snRNAseq) and spatial transcriptomics (ST); after QC, we retained 78 snRNAseq tissue samples from all 28 donors for discovery, and 43 ST tissue samples from a subset of 13 donors for replication (Fig 1C and 1D). From the snRNAseq data, we identified major cell types across all three regions from all donors, including oligodendrocytes, oligodendrocyte precursor cells (OPCs), astrocytes, ependymal cells, microglia, macrophages, vascular cells, glutamatergic neurons and GABAergic neurons (Fig 1E, 1F and Table S1). As expected, the distribution of these major cell classes varies among the three brain regions, with oligodendrocytes composing the majority of white matter nuclei, and glutamatergic neurons being restricted mostly to the DLPFC (Fig 1G and 1H). To confirm the robustness of these cell type signatures, we mapped their inferred spatial distributions in the DLPFC tissue sections ST spots (each of which contains one or more nuclei) using the cell2location^9^ workflow. Given that previous studies have localized specific glutamatergic subtypes to specific layers^10,11^, we used this mapping as a confirmation of the quality of our snRNA-seq and ST data sets. We observed oligodendrocyte and CD44-high astrocyte enrichment in the white matter adjacent to cortical gray matter (Fig. 1I), and also confirmed that our cortical glutamatergic neuron subsets (Table S2) localized to grey matter layers predicted by Azimuth human motor cortex reference^12,13^ (Fig 1J and 1K).

**Figure 1.**
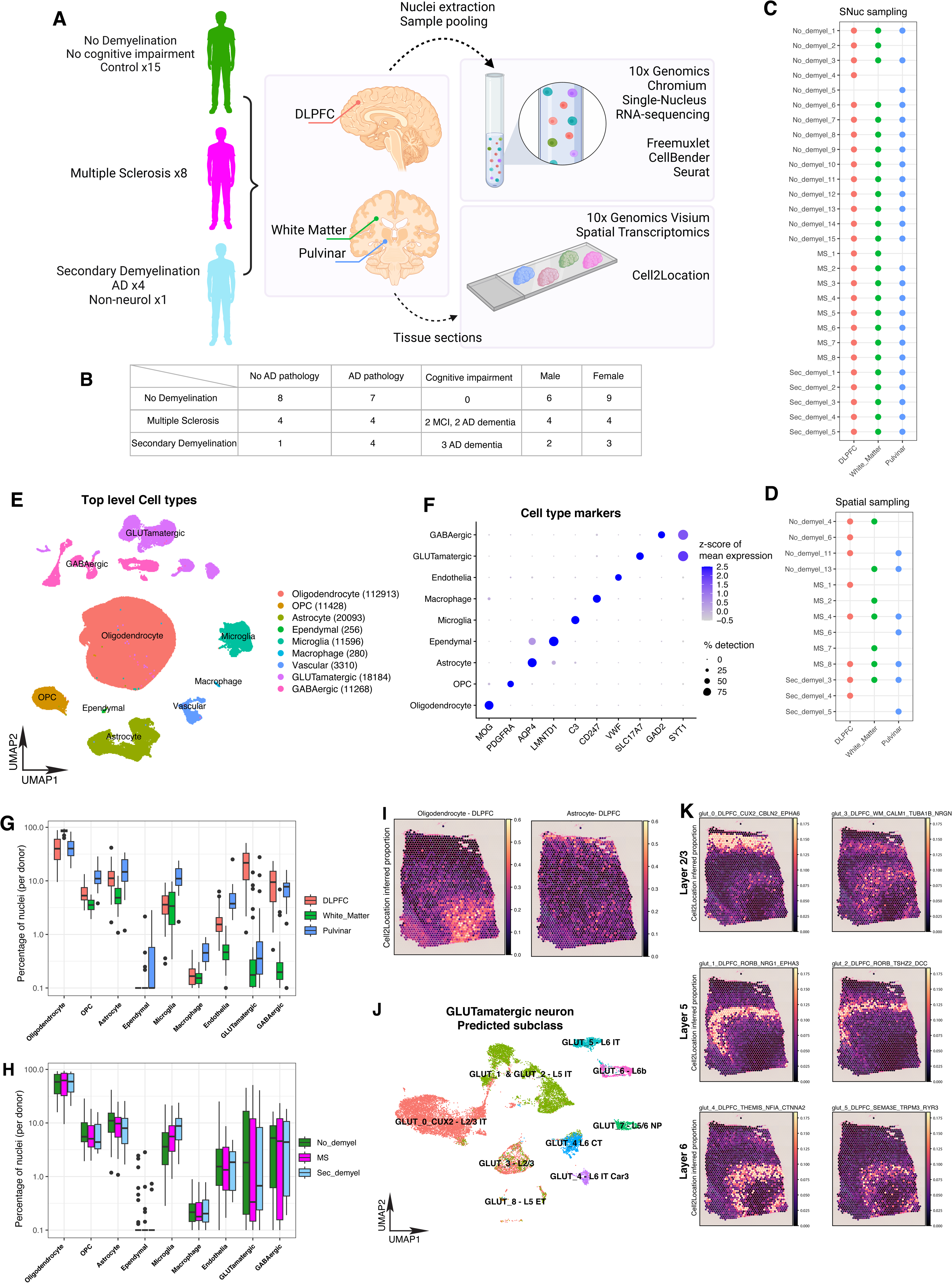
Presentation of study design, data modalities, tissue sampling, broad cell type analysis, and concordance between snRNAseq and ST data A. Illustration of study design, overview of demyelination categories, brain tissue region sampling, sequencing data modalities and applied computational methods. B. Overview of demographic, gender, cognitive impairment and AD pathology characterization across the control and demyelination groups. C. Overview of snRNAseq brain tissue and region sampling for individual donors. D. Overview of Spatial Transcriptomics brain tissue and region sampling for individual donors. E. UMAP plot for top level cell types identified in snRNAseq across three brain regions, showing segregation of major classes of cells. Numbers in parentheses indicate the total post-QC nuclei retained in the data set for each major cell class. F. Dotplot showing expression of some canonical class-specific markers in the major cell clusters. G. Boxplots showing the snRNAseq-derived proportion of each major cluster per donor across the three regions. Boxes represent the 25^th^ percentile, median (solid line), and 75^th^ percentile, and whiskers extend to the most extreme data point within 1.5 interquartile range. Individual dots represent points > 1.5x the interquartile range. As expected, there are noticeable differences in the neuronal and oligodendrocyte composition between the three regoins. H. Same as G, but with samples grouped by donor condition instead of by region. I. Representative panel showing the Cell2Location-predicted proportion of oligodendrocytes (left) and astrocytes (right) in each spot from a Visium ST experiment on a section of DLPFC tissue from a Control donor. J. UMAP plot of snRNAseq data from GLUTamatergic neurons, annotated using cortical layer labels derived from Azimuth mapping to the human primary motor cortex data from the Allen Institute. K. Same as I, but showing the predicted proportions for individual GLUTamatergic neuronal subgroups, together with their Azimuth-derived layer annotations, showing concordance between the snRNAseq and ST data for neuronal subgroups.

### A GABAergic neuronal subtype is depleted in demyelination

A central question in non-lesional tissue is the extent to which neuronal subtypes show selectively lower proportions in MS, suggestive of preferential vulnerability. In Alzheimer’s Disease, there have been reports of selective vulnerability in subtypes of deep layer pyramidal and SST+ neurons^11,14,15^. In donors with primary or secondary demyelination, however, we found only the GABA_6_CCK_LAMP5 subtype lower in the DLPFC in donors with primary demyelination and secondary demyelination (Fig. 2A, 2B Table S3). We confirmed that this effect is also found in tissue from the same individual donors with secondary demyelination using spatial transcriptomics (Fig. 2C, 2D). This distinction suggests that neuronal vulnerability in MS may be distinct from that reported in AD.

**Figure 2.**
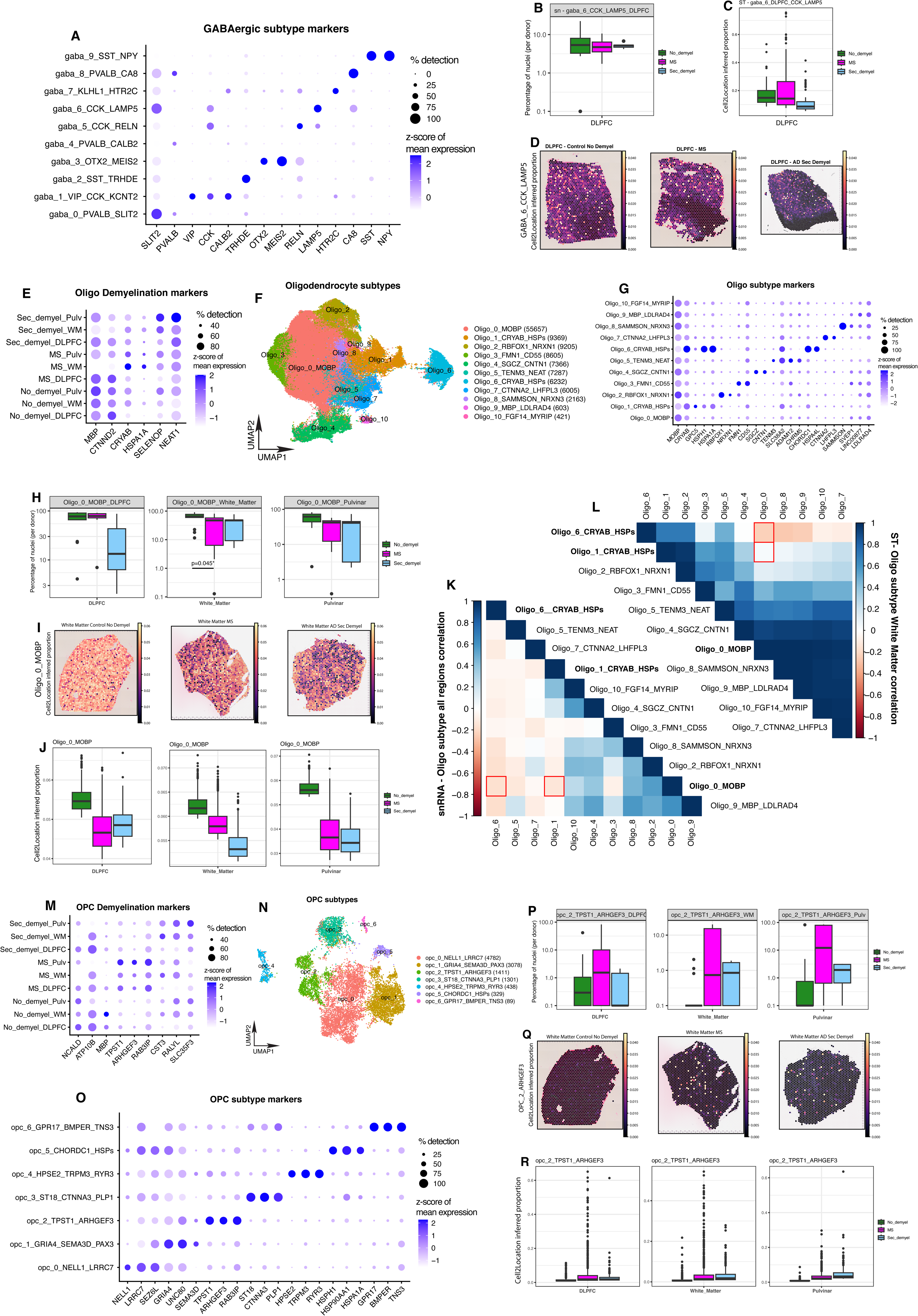
GABAergic, Oligodendrocyte and OPC subgroup composition across demyelination states, cell subtypes and brain regions. A. Gene expression dotplot highlighting marker genes for snRNAseq-derived GABAergic neuron subgroups. B. Boxplots showing the snRNAseq-derived proportion of GABAergic neuron subgroups GABA_6_CCK_LAMP5 (as a fraction of total GABAergic neurons per sample) in DLPFC across demyelination states. Boxes represent the 25^th^ percentile, median (solid line), and 75^th^ percentile, and whiskers extend to the most extreme data point within 1.5 interquartile range. Individual dots represent points > 1.5x the interquartile range. C. Same as B, but representing Cell2Location-derived proportions of GABA_6_CCK_LAMP5 per ST spot. in DLPFC. D. ST Cell2Location-derived proportions of GABA_6_CCK_LAMP5 in representative DLPFC sections from donors in each of the three groups (control, MS, and secondary demyelination). E. Dotplot showing expression of global differential genes in oligodendrocytes across donor groups in each brain region. F. UMAP plot of all oligodendrocyte nuclei post-QC, colored by cluster. Numbers in parentheses indicate how many nuclei were retained in each cluster. G. Same as A, but for oligodendrocyte subgroups. H. Boxplots showing snRNAseq-derived proportions per sample across brain regions and demyelination condition for the Oligo_0_MOBP subgroup. I. Visualization of Cell2Location-derived proportions of the Oligo_0_MOBP subgroup per spot from ST experiments on representative White Matter tissue samples across demyelination states. J. Same as H, but showing Cell2Location-derived proportions of the Oligo_0_MOBP subgroup per spot from ST experiments tissue from samples across brain regions and demyelination states. K. Pairwise correlations of snRNAseq-derived proportions of Oligodendrocyte subgroups across all regions and demyelination states. Red boxes display negative correlation between the putatively myelinating Oligo_0_MOBP subgroup and the putatively dysregulated Oligo_1_CRYAB and OLIGO_6_CRYAB subgroups. L. Same as K, but showing pairwise correlations of Cell2Location-derived proportion estimates per spot in the ST data. M. Same as E, but for Oligodendrocyte precursor cells (OPCs). N. Same as F, but for OPCs. O. Same as G, but for OPC subgroups. P. Boxplots showing snRNAseq-derived proportions per sample across brain regions and demyelination condition for the OPC_2_TPST1_ARHGEF3 subgroup. Q. Visualization of Cell2Location-derived proportions of the OPC_2_TPST1_ARHGEF3 subgroup per spot from ST experiments on representative White Matter tissue samples across demyelination states. R. Same as P, but showing Cell2Location-derived proportions of the OPC_2_TPST1_ARHGEF3 subgroup per spot from ST experiments tissue from samples across brain regions and demyelination states.

### Oligodendrocyte and OPC signatures are dysregulated in normal appearing non-lesional white matter

Given the focus of previous snRNA-seq and ST studies on oligodendrocytes and OPCs in active MS white matter lesions^5–7^, we examined these cell types in NAWM between individuals with and without demyelination due to MS. From a global perspective, we identified genes such as Myelin basic protein (*MBP*) and Alpha-crystallin B (*CRYAB*) differentially expressed in oligodendrocytes between control and demyelination groups (Fig 2E, Table S4). We then clustered oligodendrocytes into subgroups (Fig 2F, 2G, Table S5) and examined compositional differences across brain regions and demyelination states (Table S6). We observed that snRNAseq-derived proportions of the most abundant oligodendrocyte subtype (expressing high levels of *MOBP*) – Oligo_0_MOBP – was significantly lower in NAWM from MS and secondary demyelination donor tissue as compared to controls (Fig 2H), and validated this finding using spot deconvolution analysis of our ST data (Fig 2I, 2J). Genes distinguishing this Oligo_0_MOBP compared to other oligodendrocyte subgroups showed enrichment of expression programs related to cellular morphogenesis and axonogenesis (Fig S1A, Table S7), suggesting a myelinating phenotype for this subgroup. Importantly, our results show that these putative myelinating oligodendrocytes are lower in NAWM tissue in MS, suggesting differences even in the absence of active lesions in donors.

We also examined pairwise correlations between the relative abundance of oligodendrocyte subtypes across all samples, and observed that the proportion of Oligo_0_MOBP was anti-correlated with the abundance of Oligo_1_CRYAB_HSPs and Oligo_6_CRYAB_HSPs, two subgroups expressing CRYAB and heat shock proteins (Fig 2K), reflected also in our ST data (Fig 2L). Analysis of these two subgroups showed enrichment of marker genes associated with distress or dysregulation by means of active protein folding, response to temperature stimulus and response to unfolded protein (Fig S1B, S1C, Table S8 and S9). Further examination of snRNAseq-derived abundances of Oligo_1_CRYAB and Oligo_6_CRYAB showed higher proportions in our participants with MS in all three regions, and we validated this trend in NAWM in our ST data. (Fig S1D-S1F and Fig S1G-S1I).

We next performed the same analysis on oligodendrocyte precursor cells (OPCs), where we identified subgroups and genes differentially abundant between control and demyelination groups (Fig 2M, 2N, 2O, Table S10, S11). In our MS donors, we observed higher proportions of subgroup OPC_2_TPST1_ARHGEF3 in NAWM; this subgroup showed enrichment of expression programs related to cell projection organization (Fig 2P, S1J, Table S12). Our ST data confirmed that OPC_2_TPST1_ARHGEF3 was indeed enriched in tissue from donors with MS (Fig 2Q, 2R).

### Microglial signatures show distinct differences in Controls, MS, and secondary demyelination in multiple brain regions

Next, we asked whether microglial signatures differed across our three donor groups. Taking all microglia together, we identified genes – including *P2RY12*, *FTH1*, *SPP1* and *MYO1E* – differentially expressed between control and both demyelination groups (Fig 3A, Table S13), which also distinguish subgroups of microglia identified by clustering analysis (Fig 3B, 3C, Table S14). Further examination of compositional changes across groups and tissue types revealed four microglial clusters differentially abundant in control vs. demyelination groups, but in different disease-specific patterns.

**Figure 3.**
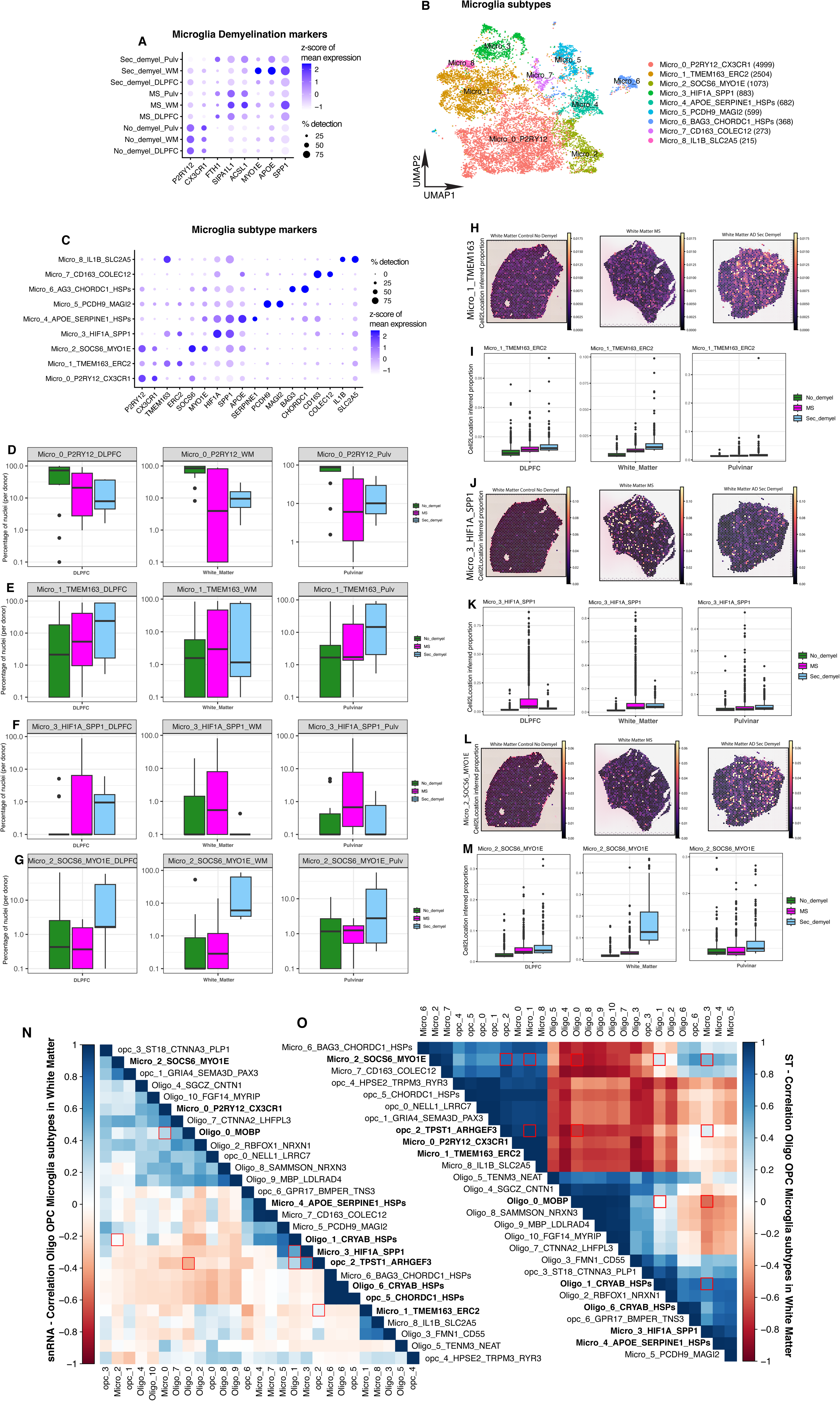
Microglia represented across cell subtypes, brain regions and demyelination states. A. Dotplot showing expression of global differential genes in microglia across donor groups in each brain region. B. UMAP plot of all microglia nuclei post-QC, colored by cluster. Numbers in parentheses indicate how many nuclei were retained in each cluster. C. Gene expression dotplot highlighting marker genes for snRNAseq-derived microglia subgroups. D. Boxplots showing snRNAseq-derived proportions per sample across brain regions and demyelination condition for the Micro_0_P2RY12_CX3CR1 subgroup. E. Same as D, but for Micro_1_TMEM163_ERC2 subgroup. F. Same as D, but for Micro_3_HIF1A_SPP1 subgroup. G. Same as D, but for Micro_2_SOCS6_MYO1E subgroup. H. Visualization of Cell2Location-derived proportions of the Micro_1_TMEM163_ERC2 subgroup per spot from ST experiments on representative White Matter tissue samples across demyelination states. I. Same as H, but showing Cell2Location-derived proportions of the Micro_1_TMEM163_ERC2 subgroup per spot from ST experiments tissue from samples across brain regions and demyelination states. J. Same as H, but for Micro_3_HIF1A_SPP1 subgroup. K. Same as I, but for Micro_3_HIF1A_SPP1 subgroup. L. Same as H, but for Micro_2_SOCS6_MYO1E subgroup. M. Same as I, but for Micro_2_SOCS6_MYO1E subgroup. N. Pairwise correlation in snRNAseq of Oligodendrocyte, OPC, Microglia subgroups across normal appearing white matter, red boxes display positive correlation between the putatively myelinating Oligo_0_MOBP and homeostatic Micro_0_P2RY12_CX3CR1 subgroup, positive correlation between the dysregulated Oligo_1_CRYAB, Micro_1_TMEM163_ERC2, Micro_2_SOCS6_MYO1E, Micro_3_HIF1A_SPP1, OPC_2_TPST1_ARHGEF3 subgroups, and negative correlation between myelinating Oligo_0_MOBP and dysregulated OPC_2_TPST1_ARHGEF3 subgroups. O. Same as N, but showing pairwise correlations of Cell2Location-derived proportion estimates per spot in the ST data.

Micro_0_P2RY12_CX3CR1 was more abundant in control donors versus donors with MS or secondary demyelination (Fig 3D), while Micro_1_TMEM163_ERC2 was more abundant in the MS and secondary demyelination groups as compared to control (Fig 3E) in all three brain regions. GO analysis of genes distinguishing these two clusters pointed to processes such as immune cell differentiation, proliferation, migration and reactive inflammatory response pathways respectively (Fig S2A, S2B and Table S15, S16). However, not all microglial signals were common to all forms of demyelination – Micro_3_HIF1A_SPP1 was more abundant only in the MS group in all 3 brain regions, and did not show the same pattern in the secondary demyelination group (Fig 3F). This suggests the existence of a microglial signature – enriched for genes associated with inflammatory activation (Fig S2C, Table S17) – that is present in non-lesional tissue specifically in MS-associated demyelination. By contrast, we observed a fourth microglial cluster, Micro_2_SOCS6_MYO1E, which showed enrichment only in the secondary demyelination group, again in all 3 brain regions (Fig 3G, Fig S2D, Table S18), suggesting that this signature may be a unique hallmark of demyelination accompanying diseases such as AD.

Given the diversity and specificity of these microglial signatures in our snRNAseq data, we interrogated them in detail in our validation ST data, and found that three of these disease-specific cell signature differences from nuclei were present in intact tissue as well (Fig 3H-3M). Overall, the condition-specific variation in microglial signatures highlights the plasticity of microglial states in different pathological conditions even in the tissue without acute demyelinating lesions, and suggests that this cell type may be responding to distributed diffusible factors.

### The NAWM demyelination signature reflects changes seen in active WM lesions

We next asked whether the gene expression and cell type differences between WM from control and NAWM from MS donors reflect those found in comparisons of WM from controls and active lesions in MS donors. We cross-referenced our data with a prior single-nucleus RNA-seq MS study that profiled cells within and at the edge of demyelinated white matter lesions at various stages of inflammation^5^, and found general agreement between our WM signatures in multiple cell types (Fig S4A, S4B). Importantly, we observed consistency in differential gene expression patterns between our NAWM samples and those from chronic active lesions across demyelination states, including genes such as *MOBP, CRYAB* in Oligodendrocytes (Fig S4C), *P2RY12*, *FTH1*, *HIF1A* and *SPP1* in Microglia (Fig S4D) and *TPST1*, *ARHGEF3* in OPCs (Fig S4E). This suggests that prior findings distinguishing control WM from active lesions are also reflected in gene expression patterns distinguishing NAWM signatures in MS donors from WM in control donors.

### Cluster correlations suggest communities of cells may change in concert in MS

Given that cell types in the brain do not act in isolation, we examined whether the proportions of cell types vary in concert, which may suggest potential co-regulation of oligodendrocyte, microglia, and OPC signatures. Identifying such “communities” of cell types may lead to further insight about putative cell-cell communication programs^11^. When examining proportions of cell type signatures from our snRNA-seq data, we found the putative myelinating Oligo_0_MOBP subgroup positively correlated with the homeostatic Micro_0_P2RY12_CX3CR1 subgroup across our donors and conditions. In addition, the MS-enriched OPC_2_TPST1_ARHGEF3 subgroup was positively correlated with the distressed/dysregulated Oligo_1_CRYAB subgroup and reactive microglia Micro_1_TMEM163_ERC2 and Micro_3_HIF1A_SPP1 subgroups, while negatively correlated with the myelinating Oligo_0_MOBP subgroup (Fig 3N), suggesting potential associations between these demyelination-associated cell signatures.

Finally, while the snRNAseq data can be used to assess communities based on cell type proportions, the ST data provides an additional axis of variation by highlighting cell signatures that are likely spatially co-localized. Here, we observed positive and negative correlations between oligodendrocyte, OPC and microglia subgroups in our WM samples. In particular, we found statistically significant co-localization of disease-associated clusters, including Oligo_1_CRYAB, Oligo_6_CRYAB, OPC_2_TPST1_ARHGEF3, Micro_2_SOCS6_MYO1E and Micro_3_HIF1A_SPP1 subgroups. Each of these clusters also appeared to be segregated from the myelinating Oligo_0_MOBP subgroup, suggesting that these disease-associated cellular signatures may be exhibit important spatial regulation in MS and secondary demyelination (Fig 3O).

## Discussion

A central question in demyelinating diseases such as MS is the extent to which non-lesional tissue is affected over the course of the disease. By examining non-lesional post-mortem human brain tissue, we found multiple glial and neuronal signatures that are altered in individuals with demyelination as compared to healthy controls. This included oligodendrocyte and OPC signatures previously reported in normal-appearing white matter, suggesting that differences between control donor WM and WM from active lesions in MS are present to a lesser degree in NAWM from individuals with demyelination. However, we cannot distinguish between the possibility that NAWM selected from our individuals previously harbored lesions and thus retain a residual signature, versus the “diffusion” of this signature globally into all NAWM from lesions elsewhere in the white matter.

A striking observation in our study is the differential proportion of specific microglial signatures in control, MS, and secondary demyelination donors. In all three brain regions we examined, HIF1A+/SPP1+ microglia are preferentially higher in MS, whereas SOCS6+/MYO1E+ microglia are higher in donors with secondary demyelination, suggesting disease-specific involvement of microglia in white and grey matter. This specificity of microglial signatures also suggests potential avenues for innate immune modulation in demyelinating diseases, with the caveat that causality between these microglial signatures and the clinical effects of demyelination require further investigation through targeted studies.

Finally, whereas these glial differences – particularly the microglial signature – are present in all regions, we find relatively fewer differences in neuronal composition among our donor groups in our two grey matter regions. The most prominent difference in neuronal composition is in a LAMP5+/CCK+ GABAergic subtype in the DLPFC, found both in the snRNA-seq data as well as validated by ST. Whereas this observation is intriguing in understanding how demyelinating diseases could affect cortical grey matter, further examination of selective neuronal vulnerability is warranted. Indeed, the lack of strong differential neuronal signals may indicate that these differences are more subtle than those observed in glia, and may be further resolved by studies with larger sample numbers, as has been noted recently in studies of Alzheimer’s Disease^11,16,17^. Alternatively, it may be the case that neuronal subpopulations do not exhibit subtype-specific vulnerability in demyelination, but rather that grey matter atrophy seen in the thalamus and the cortex is a result of global neurodegeneration affecting all cell types.

Whereas ours is the first study to to our knowledge to include both snRNA-seq and ST on a significant number of non-diseased donors and donors with MS and secondary demyelination, the primary caveat of this study is still relatively small sample size. We detect strong demyelination-associated signals in glial subtypes both in non-lesional white and grey matter, but future studies will likely uncover additional cell types with differential representation or co-localization in tissue. Despite this caveat, our study sheds additional light on the differences between NAWM and grey matter in donors with disease versus the same tissues from truly “demyelination-free” donors, and thus reveals the interplay of oligodendrocytes, OPCs, and microglia present even in non-lesional tissue in demyelinating diseases.

## Acknowledgements

This work was supported in part by grants P30AG10161, P30AG72975, R01AG15819 and R01AG17917 (ROSMAP, D.A.B.) and by R01AG066831 and R21AG075754 (V.M.). We acknowledge the data generation carried out by the Sulzberger Genome Center at Columbia University Irving Medical Center, and particularly Erin Bush and Peter Sims for establishing and running the 10x Genomics Chromium platform that generated the snRNA-seq data used in this project. This research was funded in part through the NIH/NCI Cancer Center Support Grant P30CA013696 and used the Genomics and High Throughput Screening Shared Resource. The portion of the data generation done by the Sulzberger Genome Center is also supported by the National Center for Advancing Translational Sciences, NIH, through Grant Number UL1TR001873. The content is solely the responsibility of the authors and does not necessarily represent official views of the NIH.

We also acknowledge Ryan Johnson and Gregory Klein from the Rush Alzheimer’s Disease Center for assistance in selection and provision of brain tissue from the ROS/MAP donors.

## Figure Legends

**Figure S1.**
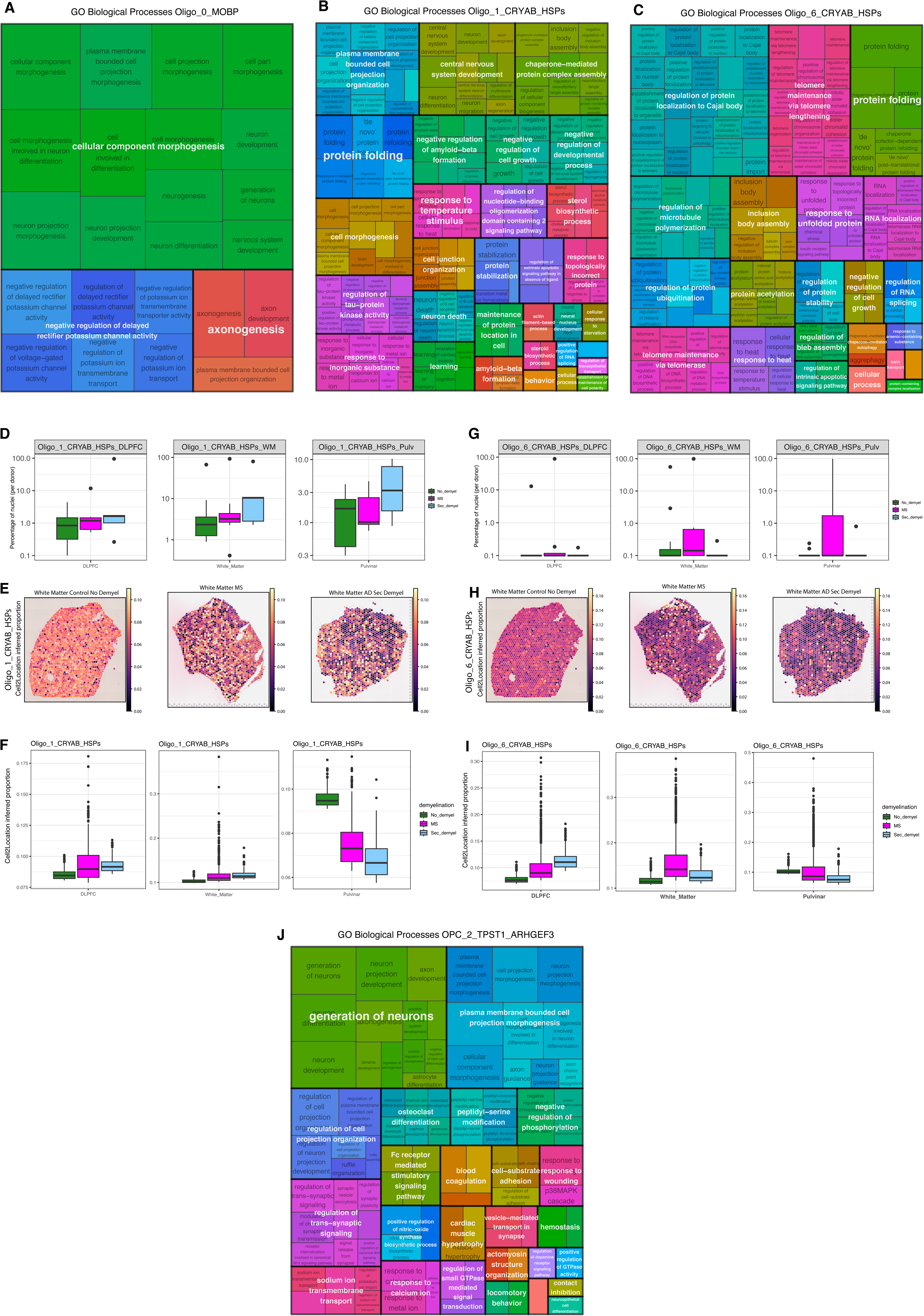
Oligodendrocyte subgroups, represented across brain regions, demyelination states. Subtype gene signatures profiled for gene ontology biological processes. A. Overview of gene ontology biological process terms in Oligo_0_MOBP for significantly upregulated genes with strong association of odds ratio above 3. B. Overview of gene ontology biological process terms in Oligo_1_CRYAB for significantly upregulated genes with strong association of odds ratio above 5. C. Overview of gene ontology biological process terms in Oligo_6_CRYAB for significantly upregulated genes with strong association of odds ratio above 3. D. Boxplots showing snRNAseq-derived proportions per sample across brain regions and demyelination condition for the Oligo_1_CRYAB subgroup. E. Visualization of Cell2Location-derived proportions of the Oligo_1_CRYAB subgroup per spot from ST experiments on representative White Matter tissue samples across demyelination states. F. Same as E, but showing Cell2Location-derived proportions of the Oligo_1_CRYAB subgroup per spot from ST experiments tissue from samples across brain regions and demyelination states. G. Boxplots showing snRNAseq-derived proportions per sample across brain regions and demyelination condition for the Oligo_6_CRYAB subgroup. H. Visualization of Cell2Location-derived proportions of the Oligo_6_CRYAB subgroup per spot from ST experiments on representative White Matter tissue samples across demyelination states. I. Same as H, but showing Cell2Location-derived proportions of the Oligo_1_CRYAB subgroup per spot from ST experiments tissue from samples across brain regions and demyelination states. J. Overview of gene ontology biological process terms in OPC_2_TPST1_ARHGEF3 for significantly upregulated genes with strong association of odds ratio above 5.

**Figure S2.**
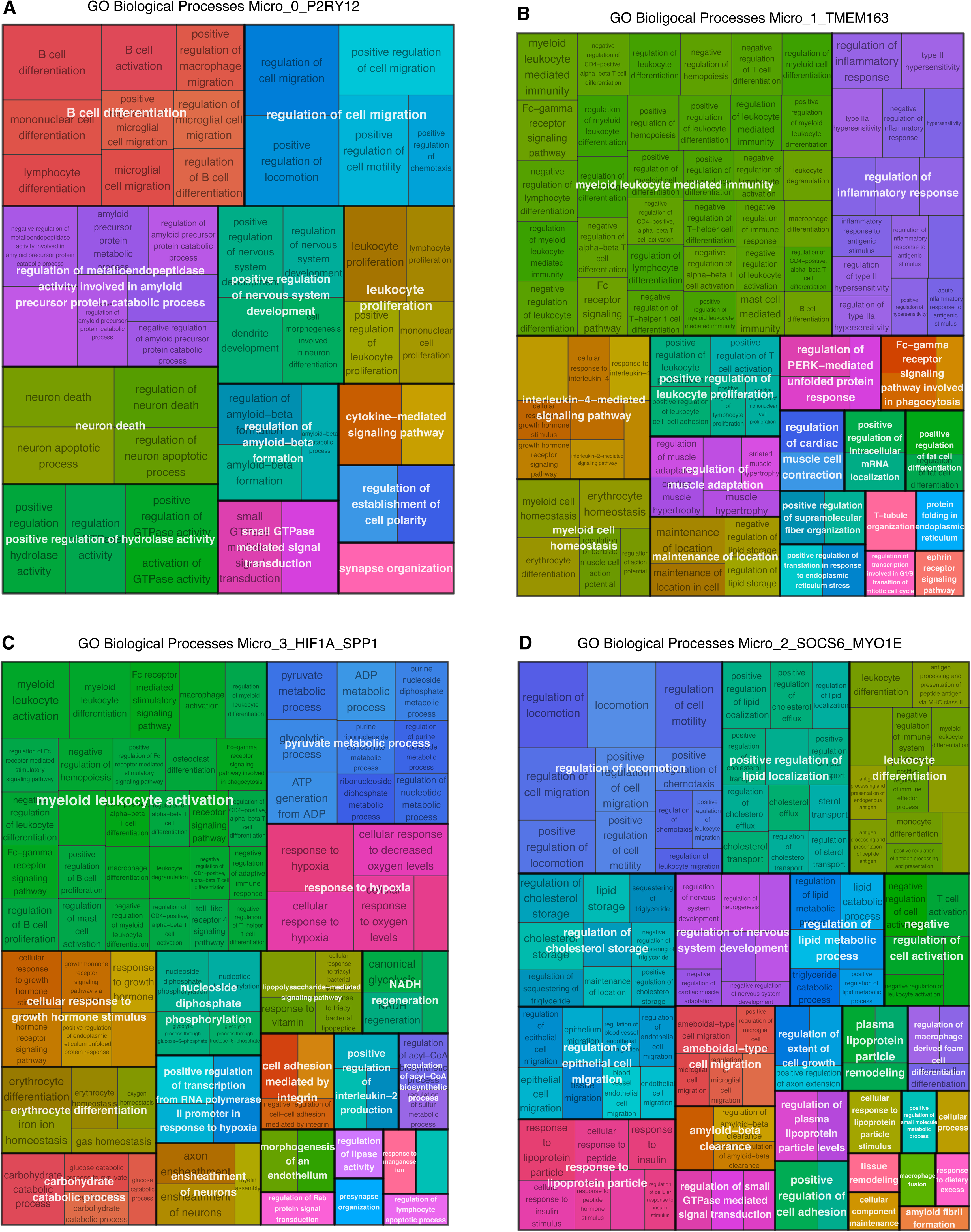
Microglia subgroup gene signatures profiled for gene ontology biological processes. A. Overview of gene ontology biological process terms in Micro_0_P2RY12_CX3CR1 for significantly upregulated genes with strong association of odds ratio above 5. B. Overview of gene ontology biological process terms in Micro_1_TMEM163_ERC2_CX3CR1 for significantly upregulated genes with strong association of odds ratio above 5. C. Overview of gene ontology biological process terms in Micro_3_HIF1A_SPP1 for significantly upregulated genes with strong association of odds ratio above 5. D. Overview of gene ontology biological process terms in Micro_2_SOCS6_MYO1E for significantly upregulated genes with strong association of odds ratio above 5.

**Figure S3.**
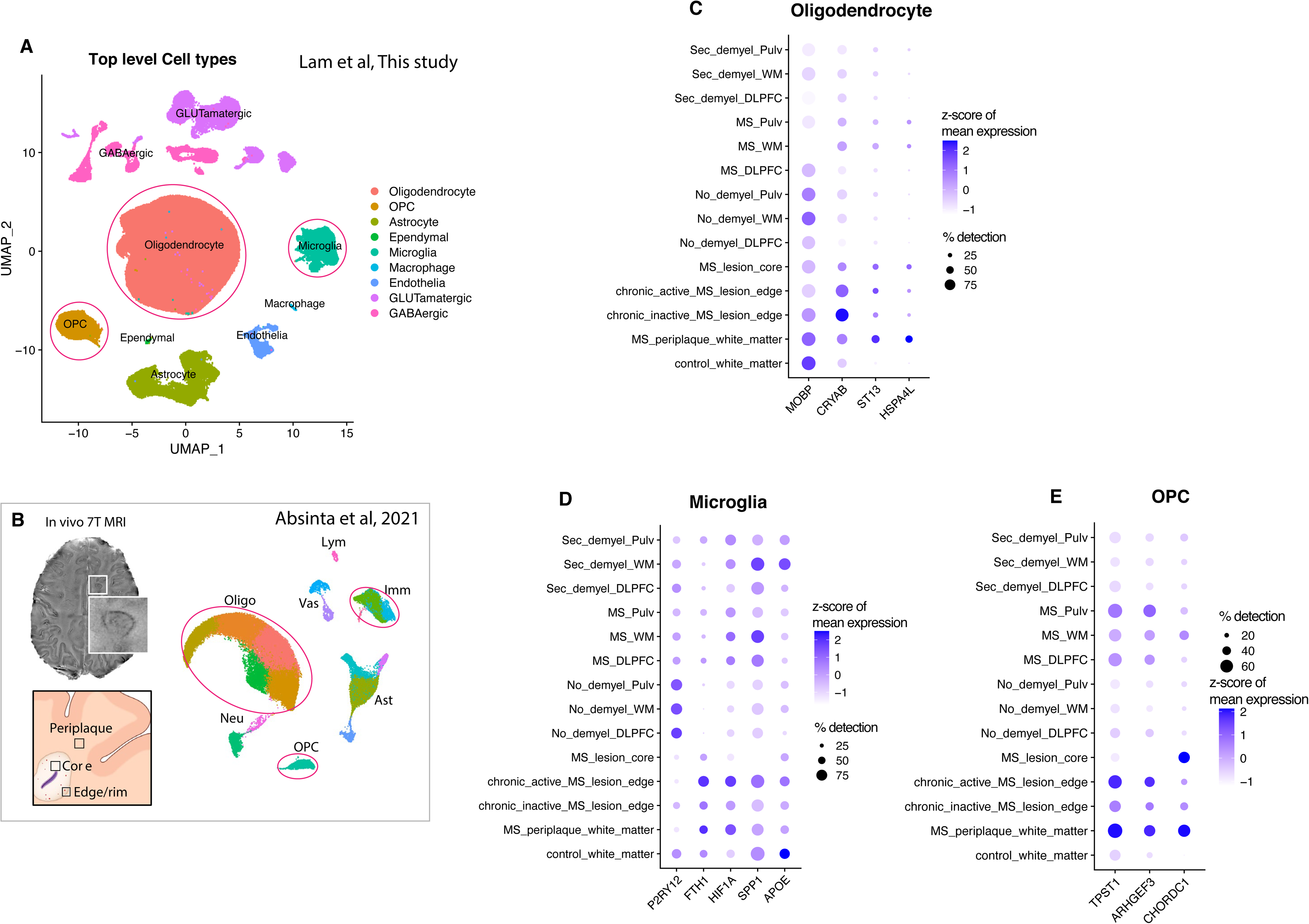
Comparing gene expression across demyelination state and region in Oligodendrocyte, Microglia and OPC toward Oligodendrocyte, immune cells and OPC in Absinta et al 2020 data set. A. snRNAseq UMAP plot overview of cell types in this data set, Oligodendrocyte, OPC, Microglia encircled. B. snRNAseq UMAP plot overview of cell types in Absinta data set, Oligodendrocyte, OPC and Immune cells encircled. C. Oligodendrocyte, dotplot of significant differentially expressed genes in demyelination states, show enriched MOBP expression in No_demyel across regions and control_white_matter (Absinta et al) and decreased MOBP expression in MS_White_Matter and chronic_active_MS_lesion_edge (Absinta et al), in contrast enriched CRYAB expression in MS_White_Matter and chronic_inactive_MS_lesion_edge (Absinta et al), enriched expression of ST13 and HSPA4L in MS_White_Matter and in MS_periplaque_white_matter (Absinta et al) D. Microglia, dotplot of significant differentially expressed genes in demyelination states, show enriched P2RY12 expression across all regions of No_demyel and in control_white_matter (Absinta et al), enriched FTH, HIF1A, SPP1 expression in MS across regions and Sec_demyel across regions and in periplaque_white_matter, chronic_inactive_MS_lesion_edge, chronic_active_MS_lesion_edge (Absinta et al), enriched APOE expression in Sec_demyel_White_Matter and in chronic_active_MS_lesion_edge (Absinta et al). E. OPC, dotplot of significant differentially expressed genes in demyelination states, show enriched TPST1 and ARHGEF3 expression across all regions of MS and in MS_periplaque_white_matter and _chronic_inactived/active_MS_lesion_edge (Absinta et al)

## Supplementary tables

Table S1. Major cell type significant gene markers.

Table S2. GLUTamatergic neuron subgroup significant gene markers.

Table S3. GABAergic neuron subgroup significant gene markers.

Table S4. Oligodendrocyte demyelination state significant gene markers.

Table S5. Oligodendrocyte subgroup significant gene markers.

Table S6. Cell subgroup differential abundance analysis significant covariates.

Table S7. Oligo_0_MOBP subgroup significant gene ontology biological processes.

Table S8. Oligo_1_CRYAB subgroup significant gene ontology biological processes.

Table S9. Oligo_6_CRYAB subgroup significant gene ontology biological processes.

Table S10. OPC demyelination state significant gene markers.

Table S11. OPC subgroup significant gene markers.

Table S12. OPC_2_TPST1_ARHGEF3 subgroup significant gene ontology biological processes.

Table S13. Microglia demyelination state significant gene markers.

Table S14. Microglia subgroup significant gene markers.

Table S15. Micro_0_P2RY12_CX3CR1 subgroup significant gene ontology biological processes.

Table S16. Micro_1_TMEM163_ERC2 subgroup significant gene ontology biological processes.

Table S17. Micro_3_HIF1A_SPP1 subgroup significant gene ontology biological processes.

Table S18. Micro_2_SOCS6_MYO1E subgroup significant gene ontology biological processes.

## Methods

### Sample description

Originally, 29 donors were selected for analysis, 1 donor was excluded due to poor quality in readout for both snRNA-seq and spatial transcriptomic, brain tissue disintegration was likely caused by inadequate preservation in the freezing process.

The 28 donors included in the analysis were as follows, 8 controls without confirmed AD pathology (ADpath) and without cognitive impairment (CTRL), 7 controls with confirmed ADpath and without cognitive impairment (CTRL_AD), these 15 controls were grouped as control without demyelination (No_demyel).

In the first disease group, 5 Multiple Sclerosis (MS) clinically diagnosed donors with microscopic confirmation of lesions following brain autopsy (Case_MS), 3 MS clinically diagnosed donors without microscopic confirmation of MS lesions following brain autopsy (Clin_MS), of these 8 MS donors 4 had confirmed ADpath. All 8 MS donors were grouped together as MS Group.

In the second disease group, 4 AD donors with confirmed ADpath (Case_AD) (3 of 4 with AD dementia) and 1 subacute infarction stroke donor termed non-neurological disease (Non_Neurol), these 5 donors were grouped together as neurological disease with demyelination (Secondary_demyel).

### Neuropathology Assessment

The mean postmortem interval was xx (SD xx). At autopsy, brains were removed and hemisected by a midline cut. One hemisphere was frozen and provided samples for this study. The other hemisphere was fixed in 4 % paraformaldehyde in 0.1 M phosphate buffer. Selected areas were processed for paraffin embedding and sectioned for neuropathology assessment of neurological diseases as described previously^8^. Plaques in multiple sclerosis cases were characterized by focal areas of white matter pallor in sections stained with hematoxylin-eosin and luxol fast blue. These areas showed preservation of axons by neurofilament (anti-neurofilament antibody, Biolegend, San Diego, CA,1:5000) immunostaining and fibrillary gliosis.

### Nucleus isolation and multiplexed single nucleus RNA library preparation

Tissue specimens from three regions, the Dorsolateral Prefrontal Cortex (DLFPC), normal appearing white matter (White_Matter) and pulvinar (Pulvinar) were received frozen from the Rush Alzheimer’s Disease Center. We observed some variability in the morphology of tissue specimens with differing amounts of gray and white matter and presence of attached meninges. Working on ice throughout, we dissected to remove white matter and/or meninges from tissue when present. We expect this to reduce variability between tissue specimens. 50-100mg of tissue per donor sample and region was transferred into the dounce homogenizer (Sigma Cat No: D8938) with 2mL of NP40 Lysis Buffer [0.1% NP40, 10mM Tris, 146mM NaCl, 1mM CaCl_2_, 21mM MgCl_2_, 40U/mL of RNase inhibitor (Takara: 2313B)]. Tissue was gently dounced while on ice 25 times with Pestle A followed by 25 times with Pestle B, then transferred to a 15mL conical tube. 3mL of PBS + 0.01% BSA (NEB B9000S) and 40U/mL of RNase inhibitor were added for a final volume of 5mL and then immediately centrifuged with a swing bucket rotor at 500g for 5 mins at 4°C. Samples were processed 2 at a time, the supernatant was removed, and the pellets were set on ice to rest while processing the remaining tissues to complete a batch of 4 samples. The nuclei pellets were then resuspended in 500ml of PBS + 0.01% BSA and 40U/mL of RNase inhibitor. Nuclei were filtered through 20um pre-separation filters (Miltenyi: 130-101-812) and counted using the Nexcelom Cellometer Vision and AO/PI stain at 1:1 dilution with cellometer cell counting chamber (Nexcelom CHT4-SD100-002). 20,000 nuclei in around 15-30ul volume were run on the 10X Single Cell RNA-Seq Platform using the Chromium Single Cell 3’ Reagent Kits v3 chemistry. Libraries were made following the manufacturer’s protocol, briefly, single nuclei were partitioned into nanoliter scale Gel Bead-In-Emulsion (GEMs) in the Chromium controller instrument where cDNA share a common 10X barcode from the bead. After reverse transcription reaction, the GEMs were broken. cDNA amplification was performed. The amplified cDNA was then separated by SPRI size selection into cDNA fractions containing mRNA derived cDNA (>300bp) which were further purified by additional rounds of SPRI selection. Sequencing libraries were generated from the mRNA cDNA fractions, which were analyzed and quantified using Qubit HS DNA assay (Thermo Fisher Scientific: Q32851) and BioAnalyzer (Agilent: 5067-4626). Libraries from 4 channels were pooled and sequenced on 1 lane of Illumina HiSeqX by The Broad Institute’s Genomics Platform, for a target coverage of around 1 million reads per channel.

### Donor and sample demultiplexing

Our single-nucleus RNA-seq libraries (PA01–PA22) used single index, which is vulnerable to index hopping. For FASTQ files of these libraries, reads affected by index hopping were detected and excluded using the algorithm proposed by Griffiths *et al*.^18^. Subsequently, the “count” command of the Cell Ranger software (v3.1.0, 10x Genomics) was used to map the reads to the human genome GRCh38 and to construct a raw UMI count matrix with custom reference transcriptome. To generate the custom reference transcriptome, a general transfer format (GTF) file of Ensembl release 91 was downloaded from the Ensembl FTP site (ftp.ensembl.org) and filtered with the Cell Ranger “mkgtf” command. To take intronic reads of pre-mRNA into account, “transcript” records in the filtered GTF file were extracted and relabeled as “exon” records. The pre-mRNA GTF file was indexed using the Cell Ranger “mkref” command. Raw UMI count matrices were processed using the CellBender^19^ software to call cells and subtract UMI counts of ambient RNA. Because nuclei from 2–4 participants were pooled together and used as input of our single-nucleus RNA-seq libraries, we executed the freemuxlet software (https://github.com/statgen/popscle) to cluster cells based on SNP genotypes of RNA reads. The “--group-list” option was specified to limit the freemuxlet analysis to cells called by CellBender. Freemuxlet generates a Variant Call Format (VCF) file for RNA genotypes of cell clusters. The RNA genotypes of cell clusters were compared with DNA genotypes of ROSMAP participants to identify origins of cell clusters. This comparison was performed using the CrosscheckFingerprints function of Picard^20^ (v2.23.6). A haplotype map for the comparison was obtained from GitHub (https://github.com/naumanjaved/fingerprint_maps).

For the DNA genotypes of ROSMAP participants, we used data of either whole genome sequencing (WGS) or genotyping arrays. The WGS genotypes (*N* = 1,196) were obtained from the Synapse website (https://www.synapse.org/#!Synapse:syn11724057), lifted over to GRCh38, and sorted by chromosomal positions using Picard. Array genotypes (*N* = 2,071) were imputed using the Michigan Imputation Server (https://imputationserver.sph.umich.edu) with the Haplotype Reference Consortium panel (v1.1) and lifted over to GRCh38 using Picard. Genotype match between RNA and DNA was assessed based on log-odds (LOD) scores of CrosscheckFingerprints. For most RNA-genotype clusters of cells, one ROSMAP participant had an outstanding LOD score compared to other participants, suggesting that these cells originated from that participant. Moreover, these outstanding participants were often in the list of pooled participants, indicating the successful demultiplexing of the cells. For some clusters, participants with outstanding LOD score were not in the list of pooled participants. In such cases, we interpreted that there was a sample swap and assigned the identity and clinical traits of the outstanding participants to the cells.

### CellBender background RNA signal removal

In each pooled sample batch run Cellranger v3 was used to map sequenced transcripts to a human reference GRCh38. CellBender^19^ was used to identify and remove background from Cellranger output “raw_feature_bc_matrix”, subsequently only nuclei designated with detected barcodes from “filtered_feature_bc_matrix” were used for single nuclei data analysis using Seurat v4.0.3^12^.

### Pre-processing of snRNA-seq data

For each sample batch run, only genes expressed in at least 10 nuclei and nuclei expressing more than 500 genes were kept for analysis. The batches were merged into one merged data object and percentage of human mitochondrial related genes were assessed; percentage of ribosomal related genes were assessed after which mitochondrial related genes were removed. Merged data object was filtered, keeping only cells with less than 5% mitochondrial related genes, more than 1.500 and less than 30.000 unique mapped transcripts.

For normalization of merged data object, Seurat function “SCTransform” was used to identify 3000 variable genes across all nuclei and variables to regress out included UMI counts, percent of mitochondrial related genes and cell cycle phase related genes. 50 principal components and 20 k-neighbors were used to resolve PC space, UMAP space and initial assignment of cell subclusters. Seurat function “FindAllMarkers” was used to find gene markers for cell subclusters using 3000 variable genes identified by “SCTransform”.

### Major cell type identification and doublet removal

In the first iteration of cell cluster assignment, major cell types were identified and manually annotated across the merged data object by gene markers identified by “FindAllMarkers” and subset into separate data objects for each cell type.

For second and consecutive rounds of quality control for assessment of cell subcluster containing doublets, in each round Seurat function “SCTransform” was used, 50 principal components and 20 k-neighbors were used to resolve PC space, UMAP space and assignment of cell subclusters. After each round Seurat function “FindAllMarkers” was used to identify gene markers for subclusters and mixed cell subclusters that contained gene markers for two or more major cell types (e.g., astrocyte/neuron, neuron/oligodendrocyte, microglia/neuron, astrocyte/oligodendrocyte, microglia/oligodendrocyte, OPC/neuron), using the following marker genes: Astrocyte (*SLC1A3*), Neuron (*SYT1*, *SLC17A7, GAD1*), Oligodendrocyte (*PLP1, MOG*), Microglia (*DOCK8*), OPC (*PDGFRA*) were removed in cell type-specific data objects.

Across the whole set of cell type-specific data objects, three to four rounds of quality control and removal of doublet cell subclusters was performed to obtain a clean readout for each major cell type.

### Subgroup identification within each major cell type class

To resolve cell subgroups for major cell types, Seurat function “FindClusters” was used to screen subclusters at range of resolutions. In order to exclude higher resolutions that split the data object into too many subclusters of similar gene expression profiles or lower resolutions that merge subclusters into clusters that showed few differences between control-disease conditions, each major cell type was assessed individually. Oligodendrocyte subtypes were resolved at resolution 0.15, Microglia subtypes were resolved at 0.25, Astrocyte subtypes were resolved at 0.15, OPC subtypes were resolved at 0.1, Endothelia cell subtypes were resolved at 0.1, Macrophage subtypes were resolved 0.25, Ependymal cell were subtypes resolved at 0.6, Glutamatergic neuron subtypes were resolved at 0.1, and GABAergic neuron subtypes were resolved at 0.1.

### Visium spatial transcriptomics sample preparation

For spatial transcriptomics, the DLPFC, white matter and pulvinar of fresh frozen brain tissue was obtained from ROSMAP cohort and dissected on dry ice. The samples from each subject were prepared into 10mmx10mm size OCT and stored in –80°C until the experiment. Samples were sectioned at 10 μm of thickness in duplicate onto a slide containing capture probes. Each tissue section contained all six cortical layers. Sections were fixed with cold 100% methanol for 30 min and then stained with H&E for 7 min at Room Temperature. Sections were scanned using a Leica Microscope. After image capture, tissue sections are permeabilized to induce cDNA synthesis and a cDNA library is generated where each molecule is tagged with its spatial barcode. The permeabilization time was optimized for the ST Visium platform and ensured that the RNA quality in fresh frozen brain tissue was sufficient for ST data generation. Only tissue sections with a RIN score above 7 were selected. The libraries were then sequenced and barcoded cDNA libraries were aligned with H&E image of the tissue section using the Space Ranger software.

### Estimation of spatial transcriptomic spot composition using Cell2location

Cell type proportions in Visium experiments were estimated using the Cell2location package in Python^9^ (version 0.1), based on the following notebook: https://cell2location.readthedocs.io/en/latest/notebooks/cell2location_tutorial.html.

Briefly, the transcriptomic signatures of each cell type were estimated from single-nucleus RNA-seq counts using a negative binomial regression. These signatures were used to estimate the relative contribution of each cell type to a given spatial capture region i.e spot in a subsequent negative binomial regression. The resulting abundance estimates were normalized in each capture region i.e spot to sum to 1.

### Differential snRNA-seq proportion analysis using ANCOMBC

Differential abundance estimation of snRNA-seq-derived proportions with respect to disease conditions was performed using the ANCOMBC^21^ R package (version 1.0.5), using demyelination status or control-vs-disease condition as the group of interest, with gender, age at death, post-mortem interval, and degree of AD pathology as covariates. A separate ANCOMBC analysis was performed for each major cell type, using only relevant cell subgroups within that major cell type. Cell subgroup differences with FDR-adjusted q-values <0.05 were then retained as being statistically significantly different across donor groups.

### Differential ST spot composition analysis

Compositional differences in cell subgroup signatures in the ST data were assessed based on the Cell2location spot-level predictions. Given the “background” levels of prediction returned by Cell2location – with very few spots containing zero values for any cell subgroups – distributions of the 90^th^ percentile of values were compared for each cell type across the disease conditions, with tissue section, donor, gender, age at death, post-mortem interval, and degree of AD pathology as covariates. Statistical significance was assessed using a Mann-Whitney test, and FDR-adjusted q-values <0.05 were retained as being statistically significantly different across donor groups.

### GO analysis to identify BP terms on significant upregulated genes in cell subgroups

For obtaining gene ontology biological process (GO BP) terms across cell subtypes, GOstats^22^ R package (version 2.64.0) was used on significant upregulated genes with fold change above 0.25 for each cell subtype. For generating GO BP term summary panel with strong association, rrvgo R package (version 1.10.0) was used on filtered GO BP terms with odds ratio above 5.

### Global differential gene expression analysis

Global differential gene expression analysis across donor conditions within each broad cell type were run using DESeq2 on pseudobulked snRNA-seq data. To generate pseudobulk RNA counts, the counts matrix was first extracted from a single-nucleus Seurat object. Counts across nuclei from the same donor, region, and broad cell type were then summed using the rowSums() function in R. The counts for each sample were then compiled into a counts table for use with DESeq2. The code used for generating pseudobulk counts is located in https://github.com/ivykos/R-Pseudobulk/

### Spatial and snRNA-seq based correlation analysis

snRNAseq counts matrices and ST count matrices for “donor region” and “cell subtype” were summarized using table() base R function to cross-classify factors and build a contingency table of the counts at each combination of factor levels. Correlation was calculated using cor() function R stats base package. Statistics and plots were generated with corrplot version 0.92 R package.

